# *S. pombe* telomerase RNA: secondary structure and flexible-scaffold function

**DOI:** 10.1101/2025.02.22.638514

**Authors:** Karen McMurdie, Allison N. Peeney, Melissa A. Mefford, Peter Baumann, David C. Zappulla

## Abstract

The telomerase RNA-protein enzyme is critical for most eukaryotes to complete genome copying by extending chromosome ends, thus solving the end-replication problem and postponing senescence. Despite the importance of the fission yeast *Schizosaccharomyces pombe* to biomedical research, very little is known about the structure of its 1212 nt telomerase RNA. We have determined the secondary structure of this large RNA, TER1, based on phylogenetics and bioinformatic modeling, as well as genetic and biochemical analyses. We find that several conserved regions of the rapidly evolving TER1 RNA are important for the ability of telomerase to maintain telomeres, based on testing truncation mutants *in vivo*, whereas, overall, many other large regions are dispensable. This is similar to budding yeast telomerase RNA, TLC1, and consistent with functioning as a flexible scaffold for the RNP. We tested if the essential three-way junction works from other locations in TER1, finding that indeed it can, supporting that it is flexibly scaffolded. Furthermore, we find that a half-sized Mini-TER1 allele, built from the catalytic core and the three-way junction, reconstitutes catalytic activity with TERT *in vitro*. Overall, we provide a secondary structure model for the large fission-yeast telomerase lncRNA based on phylogenetics and molecular-genetic testing in cells and insight into the RNP’s physical and functional organization.

## INTRODUCTION

Telomerase is a ribonucleoprotein that extends telomeres, the repeating DNA sequences that make up the ends of linear chromosomes. The telomerase core enzyme consists of a reverse transcriptase component (TERT) (1), which provides the bulk of the catalytic activity, and an RNA component (TER), which contains the template nucleotides for TERT (2). The RNA component does much more than simply provide a template for reverse transcription; it also contributes to catalysis and binds proteins involved in RNA processing, stability, and localization, as well as recruitment of telomerase to telomeres.

In the budding yeast *Saccharomyces cerevisiae*, the 1157 nt telomerase RNA (TLC1) has been shown to serve as a flexible scaffold for holoenzymatic protein subunits (3–5). The term “flexible scaffold” primarily means that the position of the functional elements that are tethered to the long RNA can be changed without causing loss of function *in vivo*. This position-independent functional modularity contrasts with the many well-studied RNPs containing subunits that must be precisely positioned relative to each other to function (e.g., the ribosome). In the case of TLC1, flexible scaffolding was first demonstrated by showing that the essential Est1 subunit could be repositioned to very different locations in the large RNA while retaining its function *in vivo* (3). Similar results were then obtained upon relocating the important Ku and Sm7 subunits (4, 6). In contrast, the functional and highly conserved RNA elements in the catalytic center could not be repositioned, although parts of the catalytic core tolerated breakage of the phosphodiester backbone (5).

Consistent with flexible scaffold architecture and function, telomerase RNAs are rapidly evolving, with large differences in both sequence and size among eukaryotes. The size of the RNA is 10-fold greater in some yeasts than in ciliates, and telomerase RNA sequences have only ~7 conserved sequence regions between *Saccharomyces* and *Kluyveromyces* budding yeasts (7). Despite the extensive sequence changes observed in the bulk of telomerase RNA during evolution, there are 4 conserved structure motifs found in the catalytic core of almost all telomerase RNAs (5, 8): the core-enclosing helix (CEH), template-boundary element (TBE), template, and pseudoknot (PK) (8). These motifs are found in this order from 5’ to 3’ and coordinate catalysis with TERT, being linked together through an Area of Required Connectivity, or ARC, so named since breaking the RNA backbone through this region also disrupts catalytic function in *S. cerevisiae* telomerase (5).

While telomerase RNAs from human and yeasts share conserved core structure motifs, they vary greatly in size, with the human RNA (hTR) being 451 nt (9) and most yeast telomerase RNAs being over 1000 nt (10–12). Yeast telomerase RNAs tend to have long, mostly helical arms connecting important structural elements, while human telomerase RNA is more compact (13, 14), yet still contains sequence-hypervariable, 3D-disordered regions (15). One of the biggest differences between hTR and the telomerase RNA from *S. cerevisiae*, TLC1, is the requirement of a three-way junction *in vitro* and *in vivo* (CR4/5 in hTR). This element is essential for catalysis with TERT in humans (16), but not in S. cerevisiae (17).

Although they are both fungi, the fission yeast *Schizosaccharomyces pombe* is evolutionarily very distant from budding yeast *Saccharomyces cerevisiae*, having diverged several hundred million years ago (18). Given this evolutionary distance and the universally rapid evolution of telomerase RNA, the sequence of *S*.*c. TLC1* was not useful in identifying *S*.*p. TER1*, but rather it was found in the *pombe* genome by inferring the template sequence from telomeric DNA repeats (10, 12). Nevertheless, the *S. pombe* telomerase RNA, TER1 (10, 12), like *S. cerevisiae* TLC1, is a very large, rapidly evolving RNA. Furthermore, initial computational lowest-free-energy predictions of its secondary structure (12) suggested it may have overall similarities to the phylogenetically and experimentally tested secondary structure of TLC1 from *S. cerevisiae* (3, 19), which is highly extended, with arms protruding from a central catalytic core. As in TLC1, TER1’s core contains the template adjacent to a template-boundary element helix (10, 12), as well as a pseudoknot (20, 21). The same groups that identified TER1 also showed that it binds the catalytic TERT protein subunit, Trt1, as well as Est1 (10, 12), homologous to the first-discovered telomerase subunit, <u>E</u>ver-<u>s</u>horter <u>t</u>elomeres 1 (Est1), from *S. cerevisiae* (22, 23). Importantly, *S. pombe* telomerase RNA 3’-end biogenesis occurs via the Sm7 complex binding to TER1 initially and then being replaced by Lsm2–8 as the RNA is processed (12, 24). Previous experiments have not shown evidence of TER1 binding the Ku heterodimer (12) or Pop 1/6/7 (23); these protein subunits bind to TLC1 in *S. cerevisiae* (25, 26) with important functions, illustrating the vast evolutionary distance and rapid evolution of the telomerase RNP holoenzyme.

As in humans and budding yeasts, fission-yeast *S. pombe* TER1 also contains a three-way junction (TWJ). This element is essential in *S. pombe* and can act *in trans* in an *in-vitro* activity assay (21, 27). The fact that TER1 has an essential TWJ makes it more catalytically similar to mammalian telomerase RNAs (16, 28) than to budding yeast, which has a dispensable TWJ (17, 29). In vertebrates, along with the catalytic core region of the RNA, the TWJ (or CR4/5 domain) also binds to TERT (16).

While a few individual components of fission yeast telomerase RNA have been studied, the only model of the overall secondary structure of the RNA published so far (12) is based solely on secondary-structure prediction software that maximizes free energy of folding. Most of this purely lowest-free-energy TER1 model remains experimentally untested, and no phylogenetic data have been published to support this software’s prediction. Evolutionarily covarying nucleotides are powerful for discovering regions of an RNA that actually base-pair *in vivo* and can be used to generate biologically accurate structure predictions (30). This approach has worked well to deduce secondary structures of ciliate, vertebrate, budding-yeast, and other telomerase RNAs (3, 15, 19, 31).

Here, we report a secondary structure model for the 1212 nt fission-yeast *S. pombe* TER1 supported by both phylogenetic analyses and lowest-free-energy modeling. We employed 7 species of fission yeasts to identify covarying base pairs and conserved-sequence regions. Furthermore, to test the model, we have evaluated the functionality of many mutant *TER1* alleles. The model allowed us to create a 623 nt Micro-TER1 allele that, like wild type, reconstitutes activity *in vitro* with TERT. We also show that the essential three-way junction element of TER1 can function when repositioned on the RNA. This finding, combined with our results showing rapid evolution of TER1 structure and nearly half of the RNA being dispensable *in vivo*, demonstrate that fission-yeast telomerase RNA is a flexible scaffold for functional modules in the RNP. Thus, overall, we provide an experimentally supported model for the fission-yeast telomerase lncRNA secondary structure, as well as insights into structure-function relationships.

## RESULTS

### Determination of *S. pombe* telomerase RNA secondary structure

To generate a phylogenetically supported model for TER1, we used sequence information from 7 *Schizosaccharomyces* species, lowest-free-energy bioinformatic predictions, and published results regarding the experimentally tested template-boundary element and pseudoknot. We first combined bioinformatic free-energy-minimization structural predictions with conserved and covarying nucleotides gleaned from TER1 alignments from *Schizosaccharomyces* species *pombe, octosporus, cryophilus, osmophilus, lindneri, japonicus*, and *versatilis* (Supplementary Fig. 1). Three of these fission yeasts, *S. osmophilus, lindneri*, and *versatilis*, were just discovered and their genomes sequenced (32–34). The alignment (Supp. Fig. 1) and a phylogenetic tree (Supp. Fig. 2) show that 4 species’ *TER1* genes are more closely related to *S. pombe* (40– 45% pairwise identity) than 2 of the others, *japonicus* and *versatilis* (~26%). Deeper divergence in regions of *japonicus* and *versatilis* species’ RNAs (Supp. Fig. 1) suggests these most-rapidly evolving parts may form different structure(s) than the other 5 RNAs. Thus, we conservatively used the more globally well-aligned 5 species (Supp. Fig. 3) for the secondary structure model-building. This led to the secondary structure for *S. pombe* telomerase RNA shown in Figure 1. In total, *TER1* alignments unveiled 14 conserved sequence (CS) regions. These are shown in the alignments in Supplementary Figures 1 and 3, as well as alongside the *S. pombe* TER1 secondary structure in Figure 1. Several CS regions are predicted to pair with each other (e.g., CS1/14, CS2/11, CS5/9, CS6/8), others form hairpin loops with the 5’ and 3’ ends of the CS region forming the stem of the hairpin (e.g., CS7, CS13), and, finally, CS10 forms the pseudoknot.

**Figure 1.**
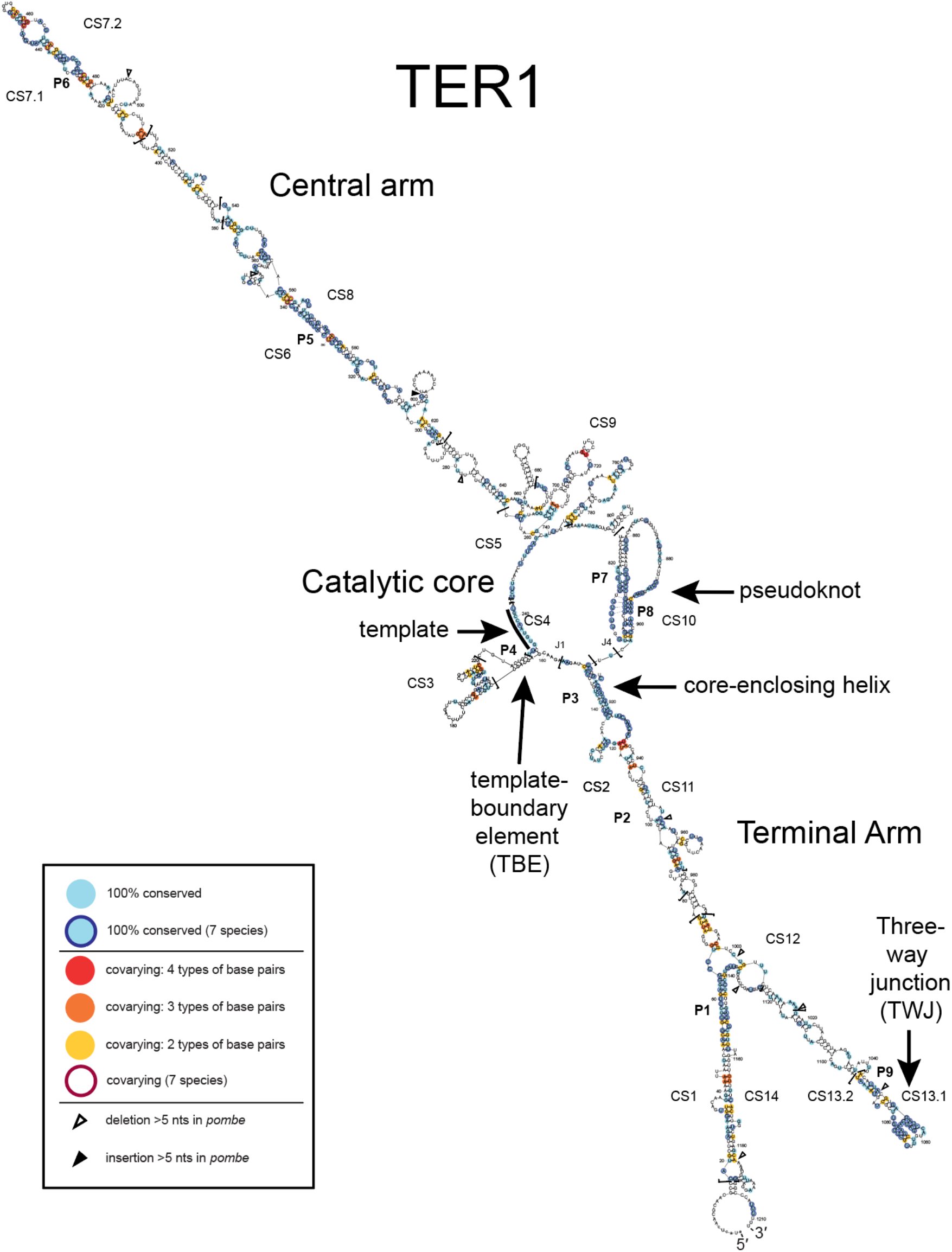
A phylogenetically supported working model of *S. pombe* telomerase RNA secondary structure. Shown is a model derived from *Mfold* lowest-free-energy predictions of TER1 constrained with evolutionarily covarying nucleotides, identified using *Alifold*. The sequences of 5 of the 7 known fission-yeast species (*S. pombe, octosporus, cryophilus, osmophilus* and *lindneri*) were aligned using *MUSCLE* (Supp. Figs. 1 and 3) and used as input for *Alifold* to help identify covarying nucleotides supporting base pairing (Supp. Table 1). The pseudoknot with base triples shown has been previously reported, as has the TBE region. Light blue nucleotides indicate 100% conservation among 5 species. Light blue nucleotides with a dark blue outline are also 100% conserved among 7 species. Other colors of nucleotides indicate that multiple types of base pairs (i.e., covarying nucleotides) are found at this location among 5 species (yellow, 2 types of base pairs; orange, 3; red, 4). Colored nucleotides with a dark maroon outline also covary among the 7 species.

We analyzed the TER1 sequence alignments of the 5 vs. 7 species using the software program *Alifold* (35), which can facilitate the identification of covarying base pairs. *Alifold* analysis yielded a rank-ordered list of covarying nucleotides that provided strong phylogenetic support for the model in Figure 1. We used 55 of the most phylogenetically supported pairings from the 5-species alignment to constrain *Mfold* predictions of otherwise lowest-free-energy secondary structures (36) (Supp. Table 1). We then manually incorporated into the model the published template-boundary element (37, 38), pseudoknot (20, 21), and single-stranded template to yield the comprehensive *S. pombe* TER1 secondary structure model shown in Figure 1.

Overall, the phylogenetic information shows that TER1 has two ~400 nt mostly helical arms that emanate from the catalytic core (Fig. 1). We have named these the Central and Terminal Arms, based on where the arms reside in the TER1 RNA transcript. The conservation data also support most of the proposed catalytic core model, such as the 3’ paired element of the pseudoknot (P8) and the core-enclosing helix (P3) (Fig. 1). We noticed that the phylogenetics also support a likely pseudoknot folding-intermediate hairpin (residues 833–906), including hairpin-specific covarying nucleotides (Supp. Fig. 4A). This newly proposed hairpin secondary structure is also consistent with reported (20) SHAPE chemical-probing data (Supp. Fig. 4B). This hairpin conformation is likely required for subsequent higher-order folding of the pseudoknot tertiary structure. A similar folding intermediate secondary structure model has been previously reported and characterized (39) for the region of *S. cerevisiae* telomerase RNA, which is in a dynamic equilibrium with the higher-order pseudoknot structure and its associated base-triple interactions.

Of the 14 conserved-sequence (CS) regions, CS5– 9 are in the Central Arm. CS6 and CS8 form a paired element, P5, in the middle of the Central Arm structure (Fig. 1). Also in the Central Arm is CS7, predicted to form the P6 paired element that coincides with the reported Est1-binding region (38). In the Terminal Arm, the three-way junction (TWJ) is formed by CS13 and is well-conserved across all 7 species (Fig. 1 and Supp. Fig. 1). In addition to the TWJ, we find 4 new CS regions in this arm, CS1, 2, 11, and 14. CS1 and 14 pair to form P1 (Fig. 1), while CS2 and 11 form P2 and P3 (the core-enclosing helix at the core). Given the conserved regions of other rapidly evolving telomerase RNAs and their determined functions (5, 26, 29, 40-43), we hypothesize that the conserved-sequence paired elements of TER1 are apt to also have RNA-folding and/or protein-binding roles in the fission-yeast telomerase RNP. **Several regions of TER1 are dispensable for function *in vivo***

Because TER1 sequence is very rapidly evolving, we hypothesized that much of the RNA would be dispensable for function. Specifically, regions where nucleotides differ greatly among species are unlikely to be forming a particular structure or have a critical function (3, 44). We used truncations in *TER1* to test the hypothesis that rapidly evolving regions would be dispensable, whereas conserved regions would be required *in vivo*. As shown in Figure 2A, we made multiple deletions in the Central (C) and Terminal (T) arms. We cultured *ter1*Δ cells harboring these truncation alleles on a plasmid for approximately 125 generations and examined telomere length by Southern blotting (Fig. 2B). We also examined TER1 RNA abundance by northern blotting (Fig. 2C) using biological replicates of cells analyzed by Southern blot in Figure 2B.

**Figure 2.**
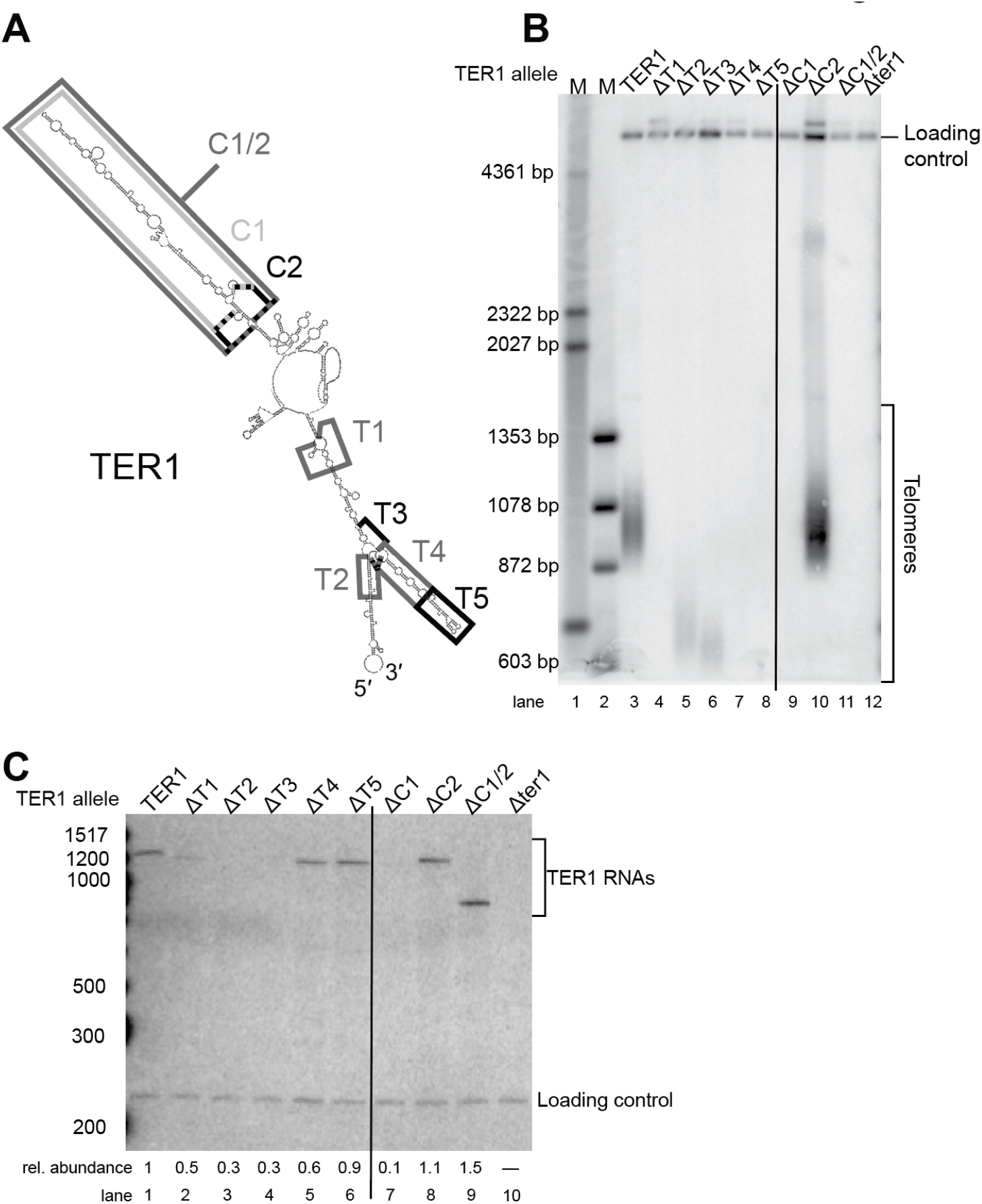
Multiple regions of TER1 RNA are dispensable for function *in vivo*. **(A)** Boxes denote regions deleted in the genetic analysis. We tested deletions based on the working model in the Central (“C”) arm as well as the Terminal (“T”) arm. **(B)** Telomeres of the *ter1*Δ strain complemented with mutant TER1 alleles and grown for approximately 125 generations. A probe was used to visualize telomeres and a probe for Rad16 was used as a loading and relative-mobility control, as performed previously. Vertical black line was added to visually differentiate between the long lanes of the blot from the single gel. **(C)** Northern-blot analysis showing the abundance of mutant alleles of TER1 RNA. Cells were grown as described in **B** and total RNA was isolated from cells before running on a 6% polyacrylamide gel. A probe for TER1 was used to identify the TER1 alleles and a probe for snR101 was used to visualize a loading control. Normalized telomerase RNA abundance was calculated by dividing the TER1 signal by the loading control signal (each with signal background subtracted in Imagequant TL software) and then the relative abundance was determined by comparing to the level for wild type. Vertical black line was added to visually differentiate between the long lanes of the blot from the single gel.

From this Southern-blot analysis, we found that ΔC2, ΔT2 and ΔT3 mutants each have stable telomeres (Fig. 2B, lanes 5, 6, and 10). Thus, regions C2, T2, and T3 are not essential for telomerase function *in vivo*. Deletion of regions T2 or T3 (but not C2) caused very short telomeres, which could be due to substantially reduced RNA abundance for these alleles (Fig. 2C, lanes 3 and 4).

We found that regions T1, T4, T5, and C1 were essential for telomerase function, as deletion caused complete loss of telomeres. The ΔT1, ΔT4, and ΔT5 alleles were nonfunctional despite appreciable levels of TER1 RNA (Fig. 2B, lanes 4, 7, and 8, and Fig. 2C lanes 2, 5, and 6). Deleting both C1 and C2 (i.e., the C1/2 allele) also caused loss of telomerase function (Fig. 2B, lane 11), consistent with the dysfunction of the ΔC1 mutant. Furthermore, 4 smaller deletions made in the C1/2 region (ΔC3, ΔC4, ΔC5, and ΔC2/3/4) were tolerated, exhibiting a shortened-telomere phenotype (Supp. Fig. 5). Thus, sub-segments of the Central Arm are dispensable for basic function but combining them to delete nearly the whole arm causes additive effects that result in a complete loss of function with respect to telomere maintenance. Other studies have shown that there are additionally important RNA sequences and RNA-binding proteins in the Est1-binding Central Arm of budding yeast telomerase RNAs, so perhaps there are also multiple components of the Central Arm of fission yeast TER1 (23, 45, 46). As for the Terminal Arm, deletion of T1, T4, or T5 does not seem to cause the observed loss of function *in vivo* because of RNA abundance, since these mutants had higher RNA levels than the fundamentally functional ΔT2 and ΔT3 alleles (Fig. 2C, lanes 2, 5, 6, compared to 3 and 4). In contrast, ΔC1 is nearly ten-fold less abundant than wild type, so a problem with RNA processing/stability/expression may be an important reason for this truncation mutant being dysfunctional (Fig. 2C, lane 7), while at the same time it is key to note that the also-dysfunctional ΔC1/2 allele did not have reduced RNA (Fig. 2C, lane 9).

### Miniaturized TER1 RNA is active *in vitro*

Since deletion analysis revealed that at least 260 nucleotides of the 1212 nt TER1 RNA are individually dispensable for function *in vivo*, we hypothesized that TER1 could be substantially miniaturized. Thus, we first designed a TER1 RNA half the size of wild-type, containing the well-conserved nucleotides in the core, as well as the three-way-junction (TWJ) region (Fig. 3A). We tested this 623 nt “Micro-TER1” RNA in a reconstituted *in-vitro* telomerase activity assay compared to full-length TER1 and an even smaller 528 nt construct that contained the core of the RNA, but not the TWJ (Fig. 3A). We synthesized these miniaturized TER1 alleles *in vitro* using T7 RNA polymerase and purified them before adding each to a transcription-translation reaction in which the Trt1 catalytic protein subunit was also produced. Each synthesized RNP was then added to a telomerase activity reaction containing an *S. pombe* telomeric primer, dATP, dCTP, and dTTP, and [α^32^P]-dGTP, as described (21).

**Figure 3.**
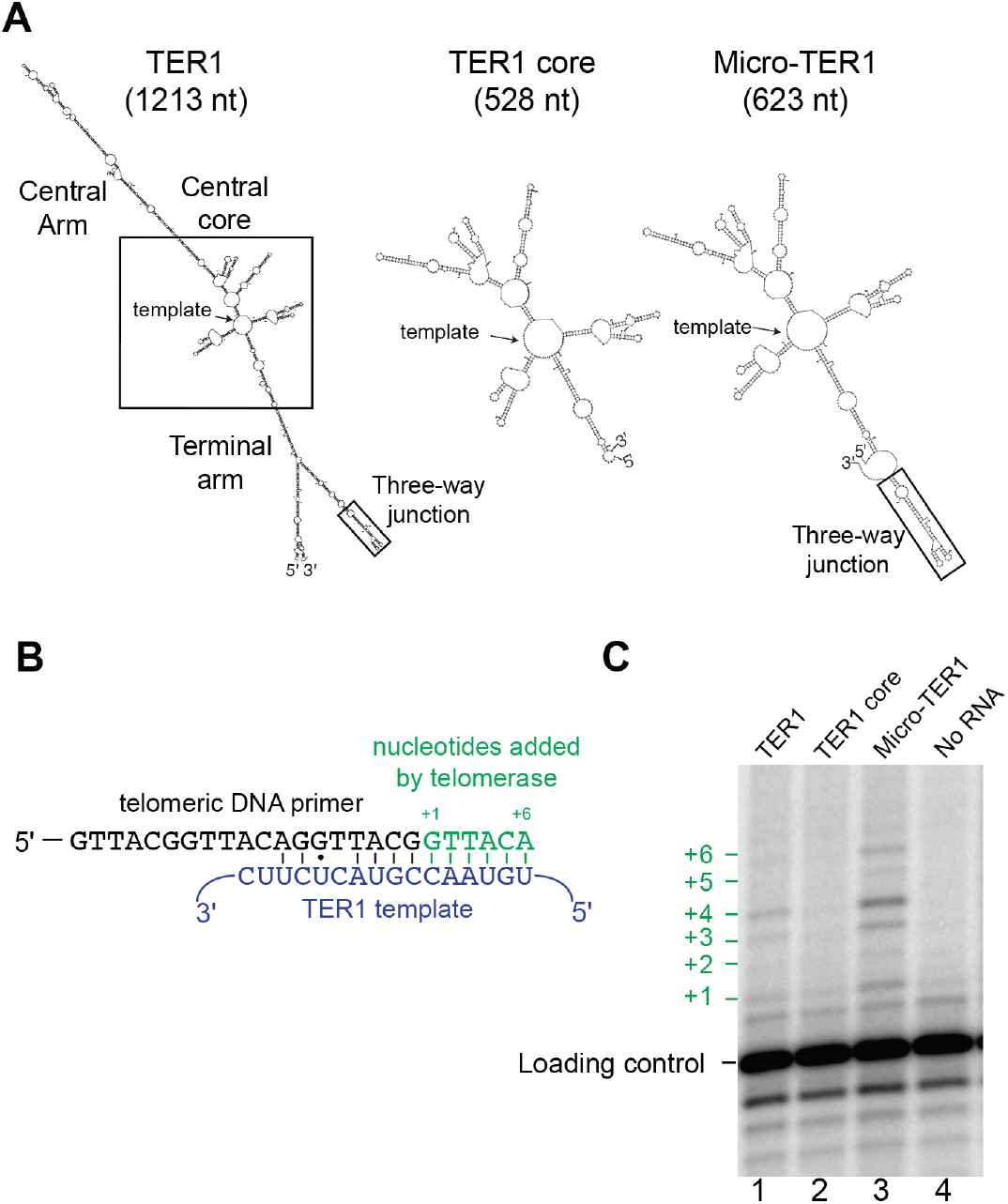
A miniaturized TER1 RNA reconstitutes telomerase activity with TERT *in vitro*. **(A)** Secondary-structure predictions of miniaturized and wild-type TER1 RNAs. The miniaturized RNAs are based on the 528-nt core region of TER1 and, independently, they still fold like the wild-type core. “TER1 core” is the 528-nt core region, and, with the TWJ added to its 3’ end, comprises Micro-TER1(623). **(B)** Schematic of the TER1 template and telomere substrate primer in the *in vitro* telomerase-activity assay. A telomeric DNA primer (black) is used as a substrate for the *in-vitro* transcribed and translated telomerase RNP complex. Blue text, the sequence of the TER1 RNA template region. Green letters, DNA nucleotides added by telomerase (the dG is radiolabeled). **(C)** Reconstituted telomerase activity radiolabeled products resolved by denaturing polyacrylamide gel. A radiolabeled template oligonucleotide was added to the activity assay to be used as a loading control. The telomeric primer used in the activity assay was not radiolabeled and is only visualized by phosphorimager analysis after the addition of alpha[32P]-labeled dGTP by telomerase.

If the *pombe* telomerase RNP enzyme is functional, it should be able to direct templated nucleotide addition to the primer starting with a radiolabeled dGTP at the +1 position and ending after the addition of 6 nucleotides (Fig. 3B). We found that the 623 nt Micro-TER1 RNA containing the three-way junction element was able to reconstitute telomerase activity, as indicated by the distinctive telomerase-specific pattern of nucleotide addition (Fig. 3C, compare wild-type control in lane 1 to Micro-TER1(623) in lane 3). On the other hand, the 528 nt “TER1 core” construct lacking the TWJ was inactive (Fig. 3C, lane 2). This is consistent with reports that the TWJ is essential for telomerase activity in both fission yeast (21) and humans (16). While all lanes have a faint band at the +1 position, it is evident from the no-RNA control that this is background signal (Fig. 3C, lane 4).

These results show that a 623 nt miniaturized TER1 RNA containing 51% of the 1212 nt mature wild-type RNA transcript is sufficient to reconstitute telomerase activity *in vitro*. The DNA primer-extension products from the telomerase enzyme containing Micro-TER1 are more intense than the bands in the full-length TER1 telomerase RNP lane, suggesting Micro-TER1 is more active than wild type. However, the amount of RNA added to the reaction was normalized by mass, and since Micro-TER1(623) is smaller than full-length TER1, the increase in intensity of DNA-product bands on the gel could be due to more telomerase complexes present in the reaction, if the RNA is limiting in the *in-vitro* RNP expression system. In summary, a half-size TER1 RNA is quite functional *in vitro*, and exhibits a pattern of nucleotide addition processivity that is similar to wild-type TER1.

### TER1 RNA serves as a flexible scaffold for the three-way junction

Given that the TWJ can function *in trans in vitro* (21) and that repositioned modules of *S. cerevisiae* telomerase RNA still function *in vivo* (3, 4, 6), we hypothesized that the presence of the TWJ is essential and positionally flexible in the large RNP. To test for such flexible-scaffold physical organization, we deleted either the local TWJ portion (T5) or the entire region (T6, encompassing T3 + T4 + T5 shown in Fig. 2A) and relocated them to two other positions on the RNA: after nucleotides 348 and 185 (Fig. 4A). To evaluate function of these novel alleles, the telomeres of *ter1*^−^ cells harboring each respective construct on a plasmid were analyzed by Southern blot after 125 generations of cell growth.

**Figure 4.**
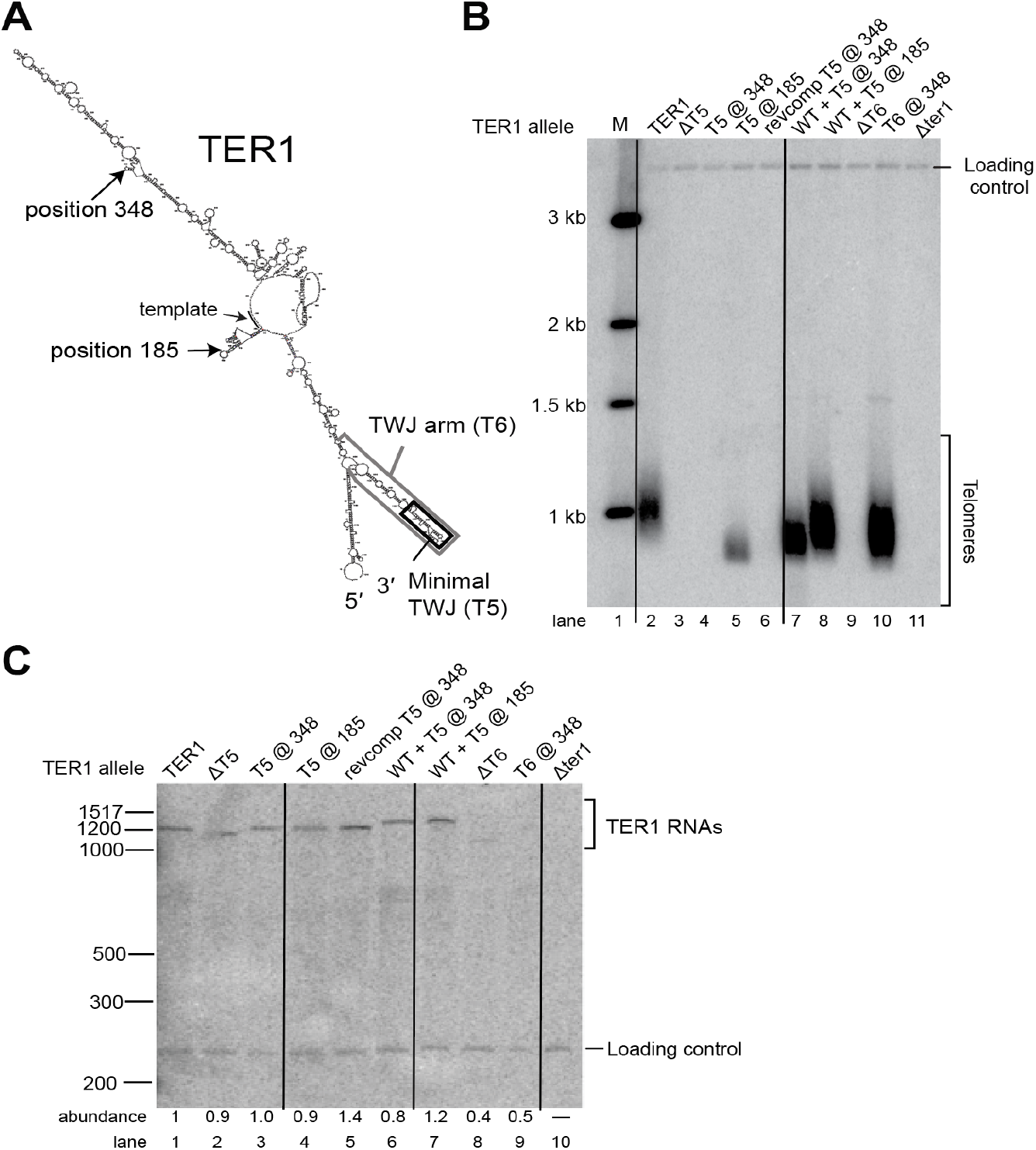
The three-way junction can function when relocated, indicating TER1 is a flexible scaffold. **(A)** Shown is the secondary-structure prediction of TER1 based on Figure 1. The three-way junction and three-way junction arm are outlined by rectangles. Sites of relocation at positions 185 and 348 are indicated by arrows. **(B)** Southern blot of telomeres from a *ter1*Δ strain expressing TER1 mutant alleles grown for approximately 125 generations. A telomeric probe was used to visualize telomeric restriction fragments and a probe for *Rad16+* was used to visualize a loading control. **(C)** Northern blot analysis of TER1 levels. Cells were grown as described in B and total RNA was purified and electrophoresed on a 6% polyacrylamide gel. Radiolabeled probes used were TER1 and, as a loading control, snR101. RNA abundance was calculated by dividing the TER1 signal by the loading control signal and normalizing to WT. Vertical black lines were added to assist as a visual aid for differentiating between gel lanes on the blot.

When the entire TWJ region (T6) was moved to position 348 (i.e., the T6@348 allele), telomeres were maintained, and at a length just slightly shorter than wild type (Fig. 4B, compare lanes 2 and 10). Similarly, the smaller TWJ region, T5, could function at another position, after nucleotide 185 (Fig. 4B, lane 5). This function depended on the inserted RNA’s structure, since the reverse-complement T5 sequence at position 185 did not support telomeres (Fig. 4B, lane 6). Telomeres of T5@185 cells were shorter than those of cells harboring the T6@348 allele (compare lanes 5 and 10). The minimal T5 three-way junction did not function at position 348, but the larger T6 region did work at this location (lanes 4 vs. 10). This may simply be due to the smaller T5 region not folding correctly in the context of insertion into TER1 at position 348 and/or it causing misfolding elsewhere. The observed phenotypes are not likely to be due to RNA abundance for these TER1 alleles, since they all accumulated to levels close to wild-type TER1, except ΔT6 and T6@348 (Fig. 4C lanes 8 and 9). However, even these two alleles with low RNA abundance were above the apparent threshold needed to maintain telomeres, as evidenced by our results in Figure 2 for the low-abundance, short-telomere-supporting ΔT2 and ΔT3 alleles. Overall, the results in Figure 4 indicate that the essential function of the TWJ can be contributed from positions hundreds of nucleotides away from the naturally occurrent location in TER1, thus showing that the large *S. pombe* telomerase RNA flexibly scaffolds the essential three-way junction element.

### Mutational analysis of the proposed pseudoknot

There have been two proposed pseudoknot models for *S. pombe* telomerase RNA in the literature (12, 21). The more recent of these two pseudoknot models (21) posits that the locations of conserved structural motifs in the catalytic conform to the overall consensus for telomerase RNAs (5, 8). The pseudoknot was shown to bind to the La protein Pof8 and a few nucleotide-substitution mutants were tailored to test these binding sites (20), yet no secondary-structure testing has been reported. Thus, to test the predicted paired elements PK1 and PK2 of the latest proposed model (Fig. 5A) (21), we used base-pair substitution and compensatory mutations predicted to disrupt and restore these hypothesized helices, respectively (Fig. 5B). If the reported pseudoknot model is accurate and its proper folding is important for telomerase function *in vivo*, we should see shorter telomeres in cells harboring alleles that disrupt pairing in these regions, while seeing telomere length enhancement in the double mutant. When we disrupted pairing by substituting nucleotides of the PK2 helix, we observed a complete loss of telomeres, indicating telomerase dysfunctionality (Fig. 5C, lanes 7 and 8). In contrast, when we combined these mutations, which is predicted to restore base pairing, we observed partial rescue of telomerase function, evident from detecting very short telomeres (Fig. 5C, lane 9; C/D compensatory mutant). These results indicate that (1) this stem within the 3’ portion of the proposed pseudoknot is essential and (2) that at least the PK2 base-pairing proposed in the model is likely accurate, given that it is striking that combining two lethal sets of base-substitution mutations improves telomerase function. The fact that the compensatory double-mutant PK2 does not function optimally could be due to the previously demonstrated critical nature of the sequence and base triples expected to be formed by this half of the pseudoknot, based on studies of budding yeasts and humans (47), as well as chemical probing of a partial pseudoknot domain of TER1 *in vitro* (20).

**Figure 5.**
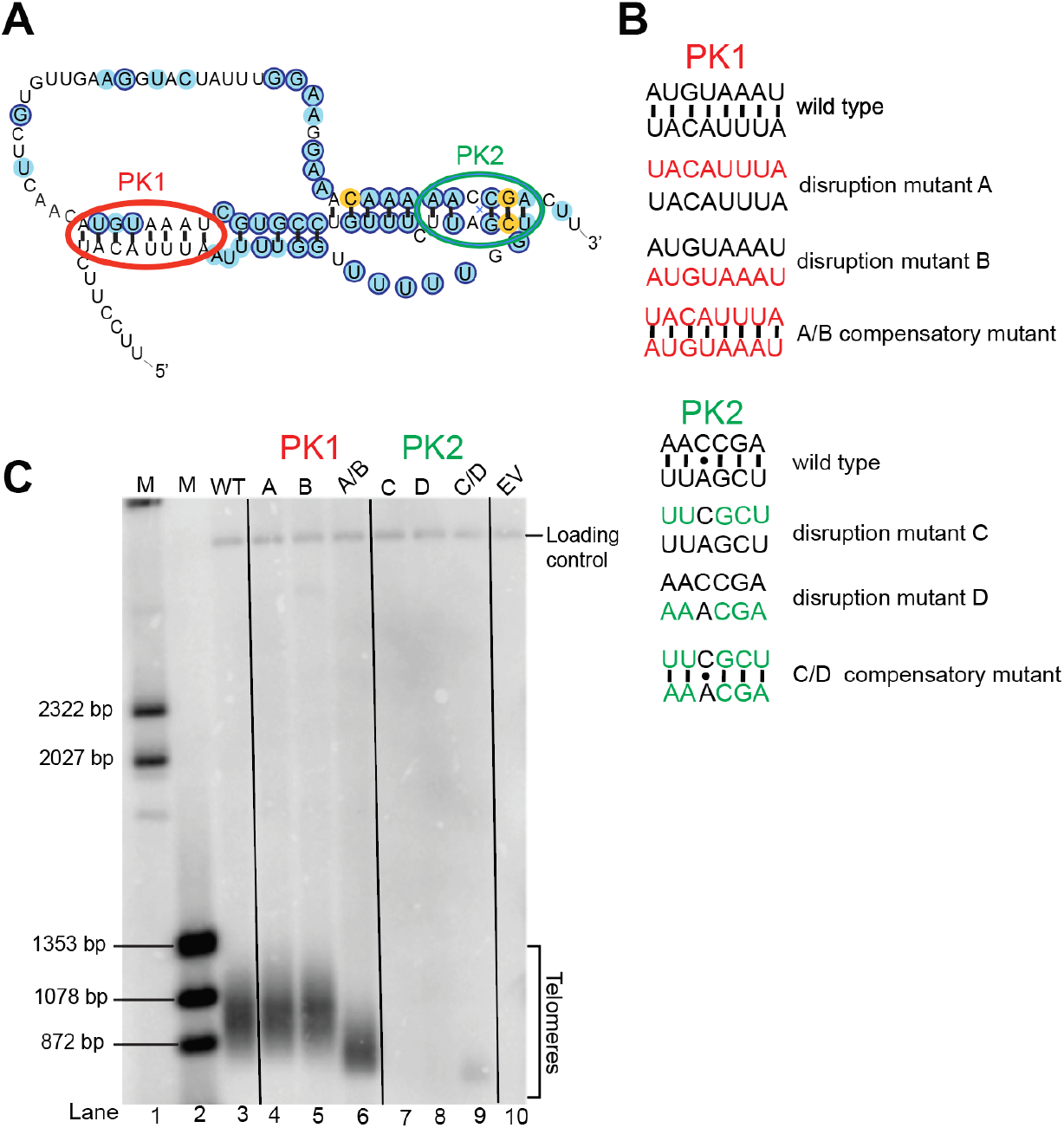
Analysis of pseudoknot model based on compensatory mutants. **(A)** A previously proposed pseudoknot model (Qi *et al*., 2012) with paired elements PK1 and PK2 circled (red and green, respectively). **(B)** Compensatory mutants used to test the pseudoknot model. Each side of each helix was mutated to its reverse-complement sequence to disrupt pairing (mutants A, B, C, and D), and then mutants were combined, predicted from the model to restore pairing (mutants A/B and C/D). **(C)** Telomere Southern blot showing the function of mutant telomerase RNAs. Cells lacking TER1 were transformed with mutant TER1 alleles and grown for ~125 generations before DNA was isolated and used for blotting. Probes for telomeres and an internal control, Rad16, were used. Vertical black lines help to visually differentiate between the long lanes.

In contrast to the results of compensatory substitution mutants in PK2, when we disrupted pairing in PK1, this did not cause any evident shortening of telomeres (Fig. 5C lanes 4 and 5). Furthermore, when we combined these sets of mutations in the same RNA, we observed telomere shortening, indicating that there is an additively negative effect of disrupting the sequence of both regions, culminating in partial loss of telomerase function (Fig. 5C, lane 6). The fact that combining the substitution mutations negatively impacts telomere length, rather than improving telomerase function, as expected if they form base pairs, suggests that the reported model (21) may not accurately reflect this stem of the pseudoknot.

## DISCUSSION

We present here the first experimentally and phylogenetically supported working secondary-structure model of fission yeast telomerase RNA, including extensive mutational tests *in vivo* and *in vitro*. At the core of the model presented in Figure 1 are highly conserved secondary structure motifs involved in catalysis: the core-enclosing helix, template-boundary element, template, and pseudoknot. We find that in TER1 these elements are in the same 5’-to-3’ orientation as other telomerase RNAs (5, 8, 49).

*Schizosaccharomyces pombe* diverged from *Saccharomyces cerevisiae* 330–420 million years ago (18). Because of this great evolutionary distance between fission and budding yeasts, when conservation is observed between these fungal species, it is often significant. This is particularly true for telomerase RNA, which, unlike other essential non-coding RNAs, is evolving extremely rapidly (3, 7, 15, 44, 48). Perhaps the most notable trait that we find to be conserved between budding and fission yeast telomerase RNA is its function as a flexible scaffold. This is the second example of a telomerase RNA acting as a flexible scaffold for the RNP’s functional elements, and the first demonstration that this kind of organizational flexibility pertains to the three-way junction element (3). Conservation of telomerase RNA acting as a flexible scaffold across

~400 million years of evolution suggests that this type of physical-functional RNP organization could be a characteristic shared with vertebrate telomerase RNAs. Support for this includes that [1] vertebrate telomerase RNAs have “hypervariable” regions in terms of both sequence and length (15), [2] the rapidly evolving portions of human telomerase RNA (hTR) have been refractory to structural characterization (13, 14), [3] hTR has been shown to be tolerant to structural perturbations, such as circular permutations, and [4] the human telomerase RNA three-way junction can function *in trans* (16, 49).

Since one of the hallmark features of flexible scaffold function of RNA in ribonucleoprotein complexes is that conserved regions are connected by long stretches of rapidly evolving sequence, which tend to be dispensable for essential function (3), we tested for this in *S. pombe*. Our analysis of *S. pombe* TER1 herein shows that at least 46% of TER1 (Supp. Table 2) can be deleted in truncation alleles (Figs. 2, 4 and Supp. Figs. 5, 6, 7), as summarized in Figure 6. Furthermore, we show that 50% of TER1 can be deleted in the Micro-TER1(623) transcript, which retains telomerase RNP activity *in vitro* (Fig. 3). These results are similar to findings regarding the large *S. cerevisiae* budding-yeast telomerase RNA, where RNAs one-third the size of wild-type (e.g., Mini-T(384); (17)) function *in vivo*, and a 170 nt RNA functions *in vitro* (5, 41). It is likely that even more of the *S. pombe* telomerase RNA is dispensable even *in vivo* than we have tested thus far. The overall secondary structure of TER1 that we report here will help guide further deletion analyses to reveal the totality of essential and dispensable nucleotides *in vivo*. Furthermore, at half the size of wild-type TER1, the Micro-TER1 RNA reported here will be advantageous for enzymological studies and can also probably be further miniaturized while supporting *in-vitro* activity with TERT.

**Figure 6.**
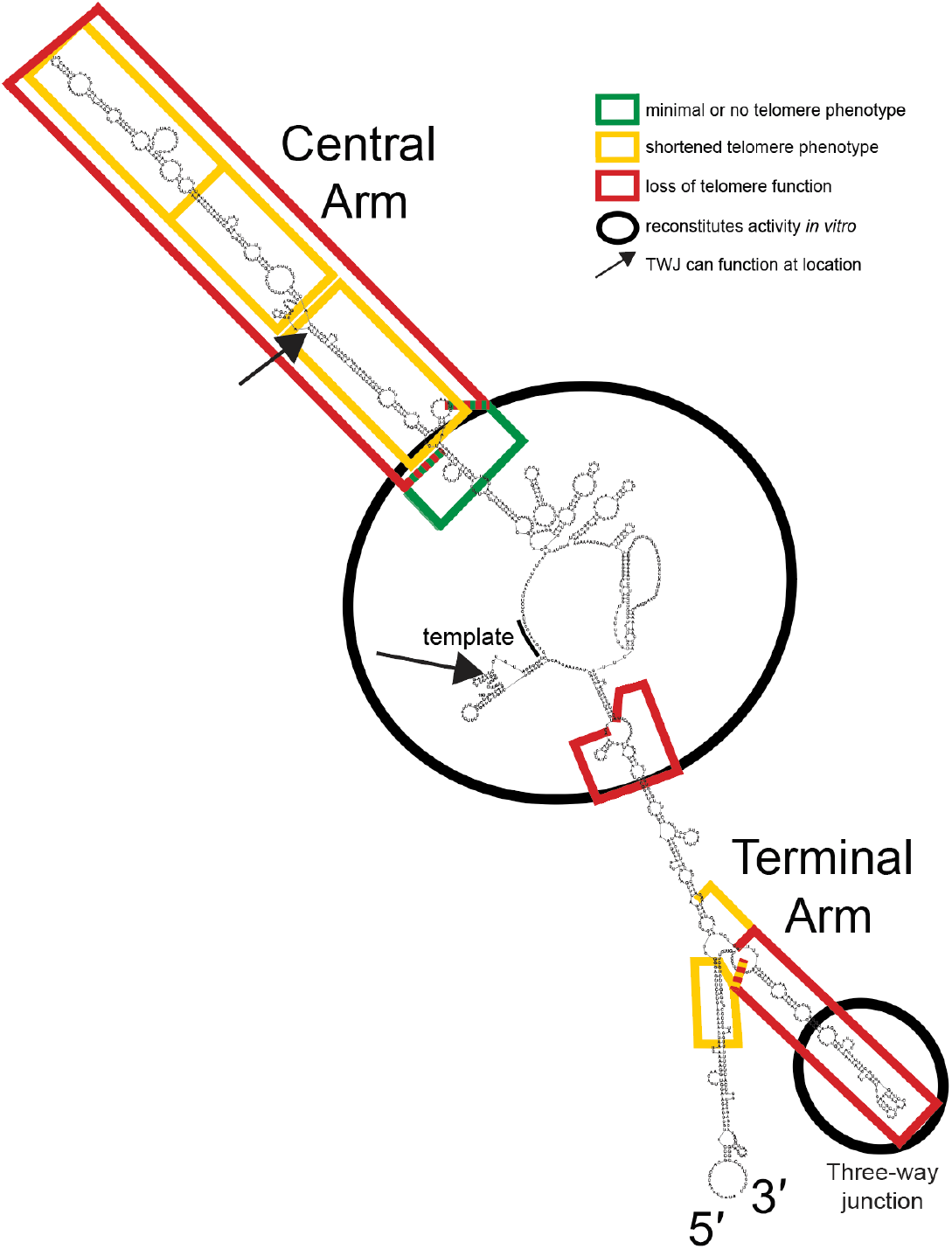
Summary of essential and non-essential regions of TER1. Regions found to be essential (red) or dispensable (yellow or green) are mapped onto a secondary-structure prediction generated with *Mfold* software. Regions where the three-way junction or three-way junction arm can function when relocated are indicated (arrows), as well as the regions sufficient to reconstitute telomerase activity *in vitro* (black circles).

Our identification of the 14 conserved-sequence (CS) regions of fission-yeast telomerase RNA should allow for the identification of all of its protein-binding sites. We anticipate that regions CS6 and CS8, which form helix P5 (Fig. 1), likely bind to a protein. The same may also be true for P1 (CS1/14) and other conserved secondary structure elements. The CS4 (template) and CS10 (pseudoknot) regions are very likely to interact with the TERT catalytic subunit, as is the CS13 (three-way-junction; TWJ) region based on what is known and demonstrated for human telomerase.

Regarding the TWJ, *S. pombe* TER1 differs from *S. cerevisiae* TLC1 in that TER1 requires it for activity *in vivo* and *in vitro* (17, 29). This makes TER1 more similar to the human telomerase RNA, which requires the three-way-junction CR4/5 domain (16). CR4/5 domain binds TERT in humans (50), so we predict that it has the same function in TER1. If the TWJ in TER1 is, in fact, binding TERT, it is likely assembled with the catalytic core of the RNA, assuming fission yeast telomerase functions as a monomer, as it does in budding yeast and humans. We have shown here that the TER1 TWJ is still able to function when repositioned to the Central Arm (position 348) or distal to the template-boundary helix (position 185), which could still allow for the TWJ to fold in 3D space to interact with the catalytic core region of the RNP.

The TER1 secondary structure model that we report here shows that the 72 nt arm extending from the template-boundary element is 253 nt shorter than the respective RNA arm in TLC1, which binds to Ku (25, 42, 51). In TER1, co-IPs have shown no evidence for Ku-binding interaction (12). In *S. cerevisiae*, the interaction between TLC1 and Ku is an additional recruitment pathway to bring TLC1 to the telomere to maintain telomere-length homeostasis (42, 51–53), so it will be interesting to see if future studies also uncover multiple telomere-recruiting pathways in fission yeast and humans.

Through our deletion analysis, we have uncovered a new essential region of TER1, the T1 region. We have shown that when T1 is deleted, telomerase is not able to extend telomeres. In an initial examination of the function of this region, we found that it does not tolerate a pair of point mutations at positions 923 and 926 (Supp. Fig. 7) and that the T1 section of the Terminal Arm must form a helix to function in maintaining telomeres *in vivo* (Supp. Fig. 6). C926 is conserved across the 7 fission yeast species, and so it may be key to TER1 function. While we still do not know what the essential function of the T1 region is, we anticipate that it is a protein-binding site. Although it is possible that the deletion of T1 could be altering folding of a structure elsewhere in TER1, *Mfold* predictions do not suggest it.

Like budding yeast TLC1, fission yeast TER1 is capable of reconstituting catalytic activity with TERT when pared down to half its normal size. In *S. cerevisiae*, however, telomerase RNA is able to function both *in vivo* and *in vitro* as 384–500-nt miniaturized forms (17), while our preliminary studies suggest that Micro-TER1 can only reconstitute telomerase activity *in vitro*. Unlike TLC1, which requires the Est1-binding region to function *in vivo*, the corresponding segment of TER1 is dispensable (38). The interaction between Est1 and TER1 functions to help bring telomerase to the telomere (38). As new studies uncover how this pathway functions in the absence of TER1’s Est1-binding region, we anticipate the ability to create a miniaturized TER1 with a truncated Central Arm that, as with corresponding “Mini-T” mutants of TLC1, can function *in vivo*.

Telomerase and telomeres are important to study for human health, since they are key to aging and age-associated diseases such as cancer, heart disease, and diabetes. It has been found that telomerase expression is activated in ~90% of human cancers (54), making this enzyme an attractive target for cancer treatments. By better understanding the structure and function of fission-yeast telomerase RNA in this genetically tractable model organism, we will be able to determine which key functional features of fungal telomerases are conserved with humans so that they can be researched more efficaciously in this advantageous model system. TER1 is only the second RNA experimentally determined to be a flexible scaffold, and we find that this type of lncRNA functional architecture is evolutionarily conserved among telomerase RNPs. This discovery advances our understanding of the generality of this non-coding RNA structure-function paradigm, further suggesting that it will apply not only to telomerase, but also to other lncRNAs.

## MATERIALS AND METHODS

### Plasmids

A full list of plasmids used is available in Supplemental Table 2. TER1 mutants tested *in vitro* were subcloned into pJW10 (10). For the *in vitro* telomerase activity assay, two plasmids containing the Trt1 gene and full-length TER1 were acquired from Julian Chen (Arizona State University) (21).

### Mutant design and construction

TER1 deletion mutants were analyzed using *Mfold* RNA secondary structure-prediction software (http://www.unafold.org/mfold/applications/rna-folding-form.php) including its *p-num* structure annotation feature to minimize off-target changes in RNA folding. Cloning strategies often involved gene-synthesized fragments (Epoch Life Science) from one unique restriction site to another in plasmid pJW10digested and subcloned into pJW10 with the equivalent region removed by restriction digest. Relocation mutants were made by deleting the nucleotides indicated and adding them to the new location. Micro-TER1 pseudoknot mutants were constructed using overlap PCR. The TER1 core construct was gene-synthesized (Epoch Life Science). Micro-TER1 was constructed by PCR-amplifying the TWJ region and adding it to the 3’ end of the TER1 core construct using an *Sph*I site. The Micro-TER1 sequence is as follows: TTAAGTATAGTTTGCTATTCCAAAACCTGTATACTGTCTACTAGAAAGAACGAGTAGTCTGTTTAGCTTTTCAGTTTGTCTGGATAGTTGTTTTCGATGGATTTCGAATTCCTGTACTGCTTCGTGTAACCGTACTCTTCAACTTTTCAGCATTGCGAAATTATTCTTTTTAGCTTTTTTTAGAGTTTTAAGCAATGAAGAAAGTTTTATAGATAGTTTCAATTCCCCATACAAATATTAAATTGTATTGGTAACAATTTTTGCTTGTGCAAATTTGTTGGCTGAATGCTCTCGTCTATACTAGCTTCTTTTGGCATTTGTTACGGAGATAAAAAGTATGGAACTTAAAAGAGCTATTATAAAAAAAATGACTTGAAGGTTTTCCTTCTACATTTAATTTTGGTTTTTTGGTCGATTCTTTGTCCGTGCTAAATGTACAACTTCGTGTTGAAGGTACTATTTGGAAGGAAACAAAAACCGACTTGTGGTCACAATGTACATTCAAACGAATAGCAACTCTGGGCATGCGGGTAAGATACTATTTTACATTACGTGAGATCCATGGATCAAAGCTTTTGCTTGTTCGCTAGTAAAATAGTTGACTACCCACCCCCCCCCCATC C

### Transformation and cell growth

Diploid strain PP407 was patched onto malt-extract medium and incubated at 28°C for 3 days to induce sporulation. Once tetrads were seen under the microscope, they were digested with glusulase overnight before plating spores onto YEA+G418 media. A single colony was selected and grown in liquid medium at 32°C, then transformed with TER1 mutant alleles using a standard electroporation protocol. Transformants were plated on minimal medium lacking uracil to select for the pJW10 plasmid and its derivatives.

Once cells were transformed with the TER1-mutant alleles, they were grown at 32°C and restreaked onto fresh plates every 3 days. Cells from restreak 3 were grown in minimal medium lacking uracil until saturated and then collected by centrifugation. This pellet was split, half was used for isolating genomic DNA and the other was used for isolating total cellular RNA.

### Southern blots

Southern blots were performed as described in reference (17), with the following exceptions. DNA for the Southern blot was digested with *Eco*RI overnight, then electrophoresed through a 1.1% agarose gel at 70W for 16.5 hours. DNA was transferred to a Hybond-N+ nylon membrane by capillary action and UV-crosslinked. The membrane was probed overnight with a telomeric probe as well as an internal control fragment from the *rad16+* gene.

### Northern blots

Northern blots were performed as described in (17) with the following exceptions. 10 μg of RNA was run on a 4% denaturing polyacrylamide gel at 35W for 3.5 hours. The RNA was then electrotransferred to Hybond-N+ nylon membrane (Cytiva) and UV-crosslinked. The membrane was probed with nucleotides 640–880 of TER1 DNA labeled with ^32^P-dCTP using random priming, as well as an end-labeled oligonucleotide

(5’-CGCTATTGTATGGGGCCTTTAGATTCTTA-3’) complementary to a region of SnR101 as an internal loading control.

### RNA synthesis *in vitro* and purification

Templates for RNA synthesis were made by PCR-amplifying the region to be transcribed using a forward primer containing the T7 promoter followed by 3 guanosine nucleotides. These templates were then used in a typical T7 *in-vitro* transcription reaction and incubated at 37°C for 1 hour. RNA was then ethanol-precipitated before being run through a 6% denaturing polyacrylamide gel for ~4 hours. The gel was stained using GelStar (*Lonza*) and RNA was visualized by ultraviolet light. The band for each RNA construct was cut from the gel, crushed, and incubated with rotation in equal parts TE (pH 8) and phenol/chloroform/isoamyl alcohol, pH 6.7, and 0.1 M sodium acetate at 4°C overnight. Gel fragments were removed from the mixture using a column made of autoclave-sterilized glass wool and glass beads. RNA was then phenol-chloroform extracted and ethanol-precipitated.

### Activity assay

The Trt1 protein was made using a rabbit reticulocyte lysate (RRL) transcription and translation system (Promega). After incubating at 30°C for 1 hour, the reaction was divided into aliquots of 10 μl and then 1 μg of previously T7-transcribed and gel-purified RNA was added to each aliquot. This mixture was then incubated at 30°C for 30 min.

To test for telomerase activity, 4 μl of the RRL + RNA mixture described above was added to a reaction containing 1X activity buffer (50 mM Tris– HCl at pH 8.0, 50 mM KCl, 1 mM MgCl2, 5 mM 2-mercaptoethanol, 1 mM spermidine), 1 μM telomeric DNA primer (PB872), 0.5 mM dATP, 0.5 mM dTTP, 0.5 mM dCTP, 2.92 μM dGTP, 0.33 μM ^32^P-dGTP (3000 Ci/mmol), and ~5000 cpm of ^32^P end-labeled PB872 as a loading control. This reaction was incubated at 30°C for 30 min, then stopped by adding 100 μl of stop buffer (10 μg/ml glycogen in 3.6 M ammonium acetate). The reaction was then ethanol-precipitated and run on a 10% acrylamide/7M urea gel at 90 W for 75 min.

### Secondary-structure modeling

A *MUSCLE* multiple-sequence alignment of *TER1* from the five most closely related species (*pombe, lindneri, cryophilus, osmophilus*, and *octosporus*) was generated and entered into *RNAalifold*. The *Alifold* base-pair output was used to identify covariation events. The top 74 covarying base pairs were used for next steps, since after this point nucleotide conservation instead of covariation began appearing in the rank-ordered list from *Alifold*. Base pairs that were isolated or that covaried in other species but were absent in *S. pombe* were removed. The remaining 55 bp identified as covarying by *Alifold* were used to constrain *Mfold* (Supplementary Table 1). For the final structure model, additional *Mfold* constraints were the 4 bp in the TBE helix (37) and the template being single-stranded. The core was manually modeled to accommodate these data as well as the published pseudoknot (20). Every base pair in the model was analyzed to determine if covariation or univariation (variation of G to A or *vice versa* to maintain pairing with a U residue) occurred in the 5-way alignment but was not in the top results from *Alifold*. Another 7-way MUSCLE alignment was generated including the two other more distant species, *S. japonicus* and *versatilis*. There were 13 covariation events identified among the 7 species. Three of these base pairs occur in the *Mfold* before the pseudoknot was manually added (Supp. Fig. 5). Six of these base pairs were localized to one region where 3 base pairs already existed in the model and 3 were slightly shifted.

## AUTHOR CONTRIBUTIONS

KEM, MAM, PB, and DCZ designed the wet-bench experiments. ANP and DCZ performed the final model-building and related analyses upon the discovery of 3 new fission yeast species in 2023 after KEM and DCZ performed initial phylogenetic analysis and modeling. KEM and MAM made the plasmid constructs, designed with DCZ, and did all *in-vivo* experiments and blots. All experiments were performed by KEM except for final version of Figures 1, Supplemental Figures 1–4 and 6. PB provided the *Schizosaccharomyces pombe* strains. The manuscript was first drafted by KEM and DCZ and later revised by DCZ and ANP.

## ACKNOWLEDGEMENTS

This research was supported in part by the National Institutes of Health (National Institute of General Medical Sciences) grants R00 GM80400 and R01 GM118757 to DCZ and funds to DCZ from Johns Hopkins University, Lehigh University, and Mike and Joan Hoben. KEM was supported in part by National Institutes of Health Cellular and Molecular Biology graduate student training grant 2T32 GM007231. This work was also supported in part by funds to PEB from the Howard Hughes Medical Institute and the Stowers Institute for Medical Research.

## Notes

### Competing Interest Statement

The authors have declared no competing interest.

