## Supplementary figures and tables for "*S. pombe* telomerase RNA: secondary structure and flexible-scaffold function"

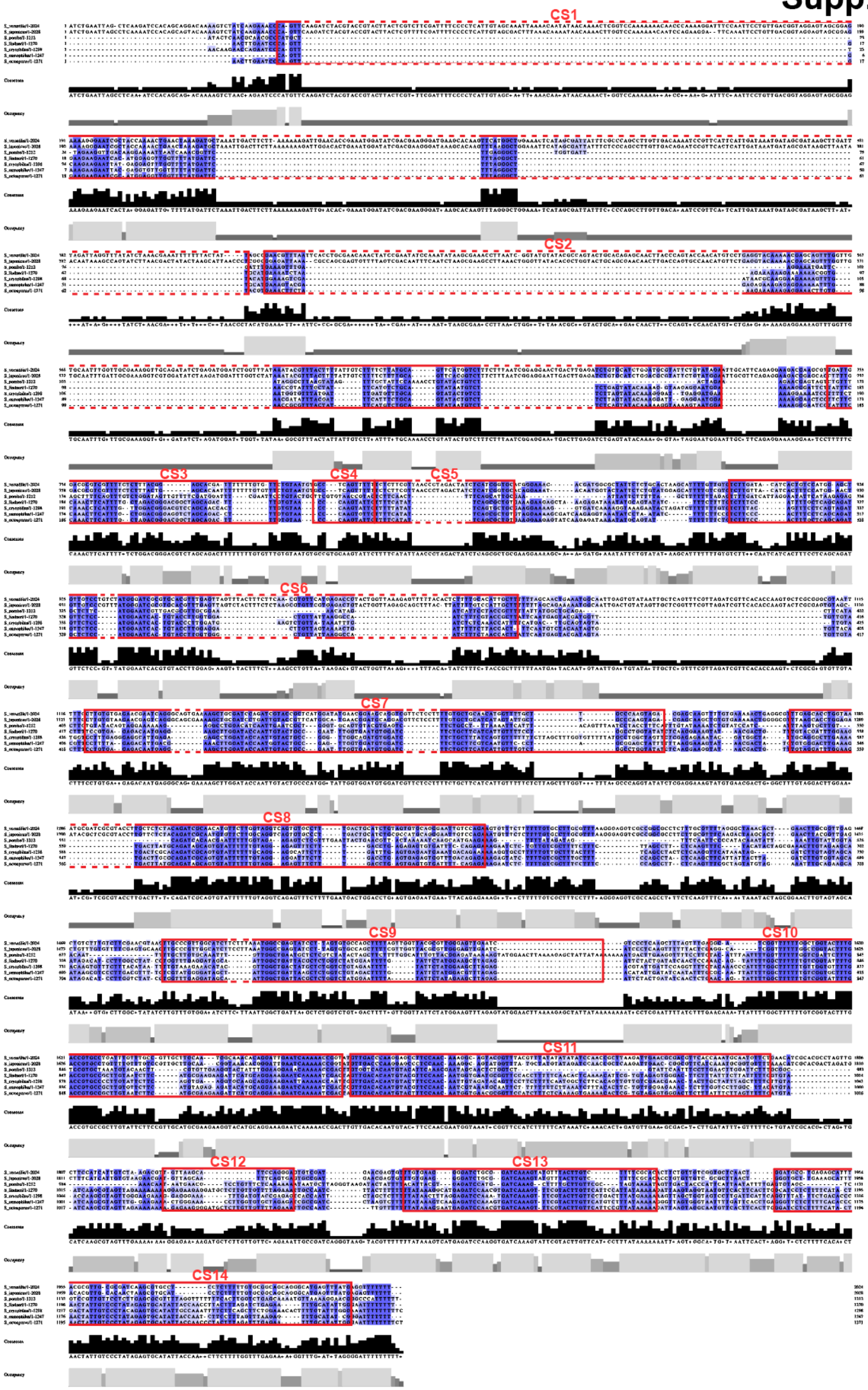


**Supplementary Figure 1. Alignment of TER1 telomerase RNA gene from 7 *Schizosaccharomyces* species.** *MUSCLE* was used to generate a multiple-sequence alignment of TER1 from the 7 species. The alignment was visualized using *Jalview* software and exported with base identities highlighted. Darkest blue color indicates 100% conserved nucleotides, with lighter tones indicating somewhat lower levels. Red boxes indicate the conserved sequence (CS) regions 1–14, which are also indicated on the TER1 secondary structure in Figure 1. A Jalview-generated consensus sequence is indicated below the aligned sequences.

#### Supp. Fig. 2

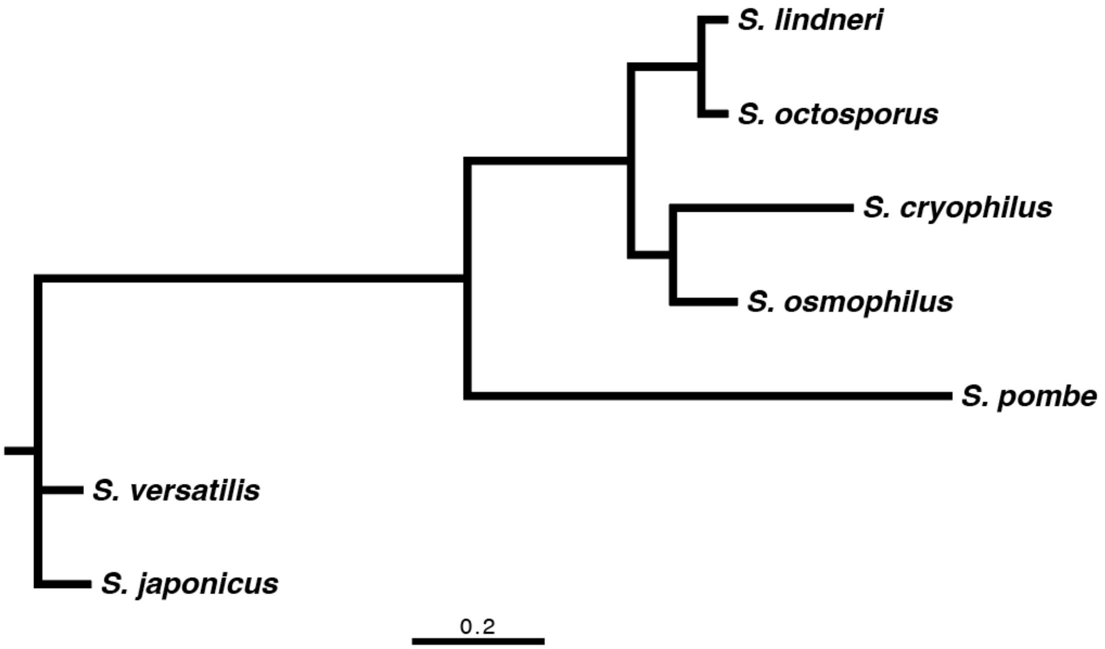


**Supplementary Figure 2. Phylogenetic tree showing relatedness of the 7 *Schizosaccharomyces* species' TER1 genes.** *IQTree* was used to generate the phylogenetic tree for TER1 based on the alignment in Figure 1, which was then visualized using *FigTree* software. The sequences of 5 species (*S. lindneri*, *octosporus*, *cryophilus*, *osmophilus*, and *pombe*) are more similar to each other than the outgroup containing *S. versatilis* and *japonicus*. Scale bar, rates of change based on *IQTree* calculations.

**Supp. Fig. 3**

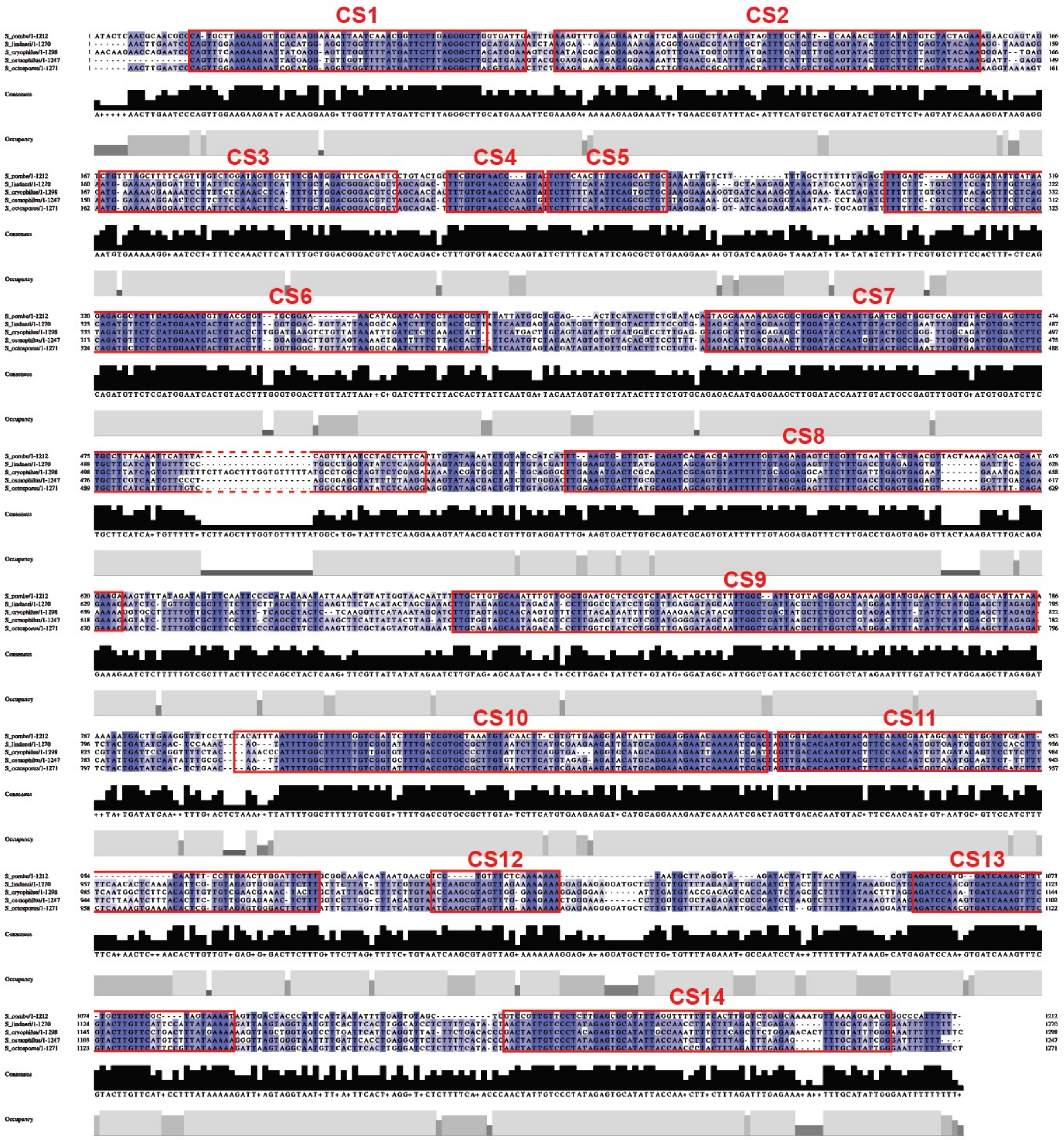


**Supplementary Figure 3. Alignment of TER1 from the *Schizosaccharomyces* species most related to *S. pombe*.** As in Supplementary Figure 1, TER1 sequences were aligned from 4 species most related to *S. pombe*, and thus 5 are shown in the alignment output, as visualized using *Jalview* software. Darker blue color indicates greatest conservation, whereas white regions are not conserved in sequence. Red boxes indicate the conserved sequence (CS) regions as shown in Figure 1 (secondary structure model of TER1) and Supplementary Figure 1 (7-species alignment). A Jalview-generated consensus sequence is indicated below the aligned sequences.

A

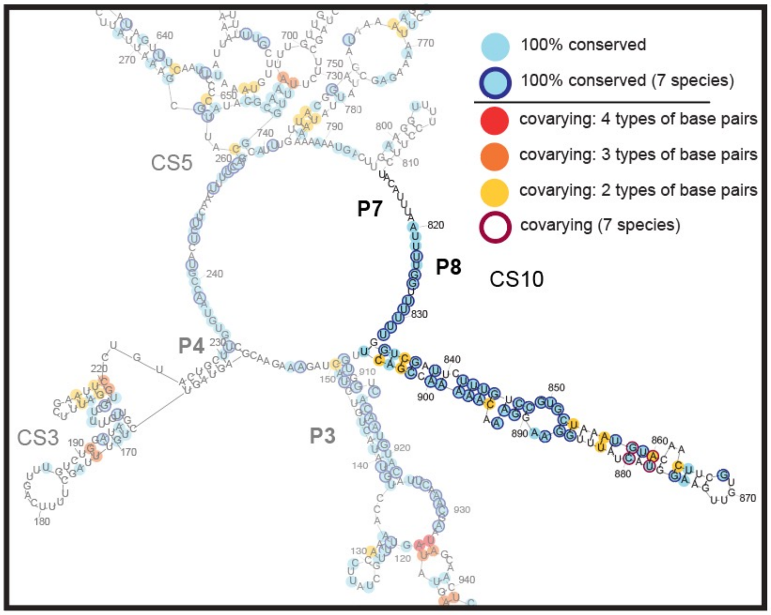


B

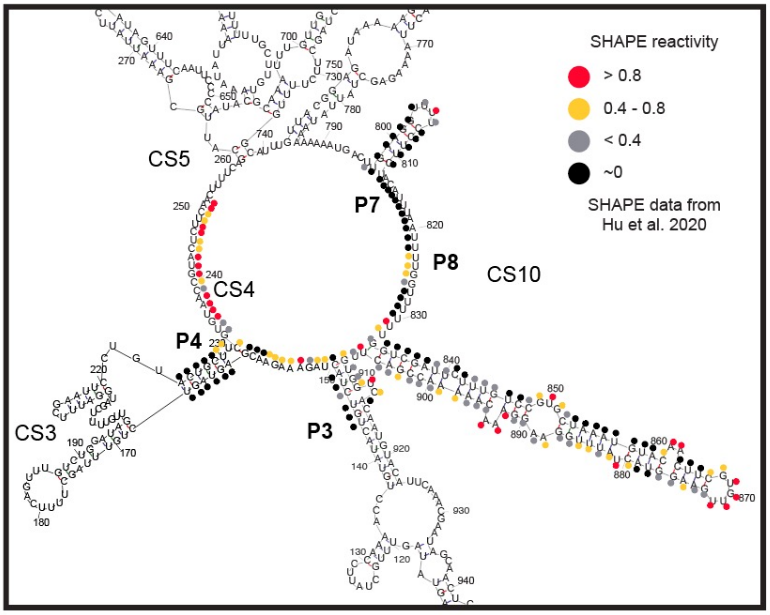


**Supplementary Figure 4.**

**Proposed secondary structure model of the core of TER1 (without the pseudoknot or base triples).** The model shown in both A and B was determined using *Mfold* (which does not predict pseudoknots) and forcing the base pairs listed in Supp. Table 1, which are supported by covarying nucleotides during evolution. The structural model is the same as Figure 1 (where the higher-order folded state including the reported pseudoknot is shown) except for the region that is not faded-out in **A**; i.e., the residues that give rise to the modeled pseudoknot.

**A. Conservation and covariation data mapped onto our secondary structure model (i.e. without the pseudoknot).** Some base pairs are separated in the pseudoknot but are predicted in the secondary structure model, which could be a folding intermediate (pseudoknots are higher-order RNA structures that fold after RNA initial base-pairing interactions in helices).

**B. SHAPE reactivity data from Hu *et al.* 2020 mapped onto our secondary structure model of the core.**

Supp. Fig. 5

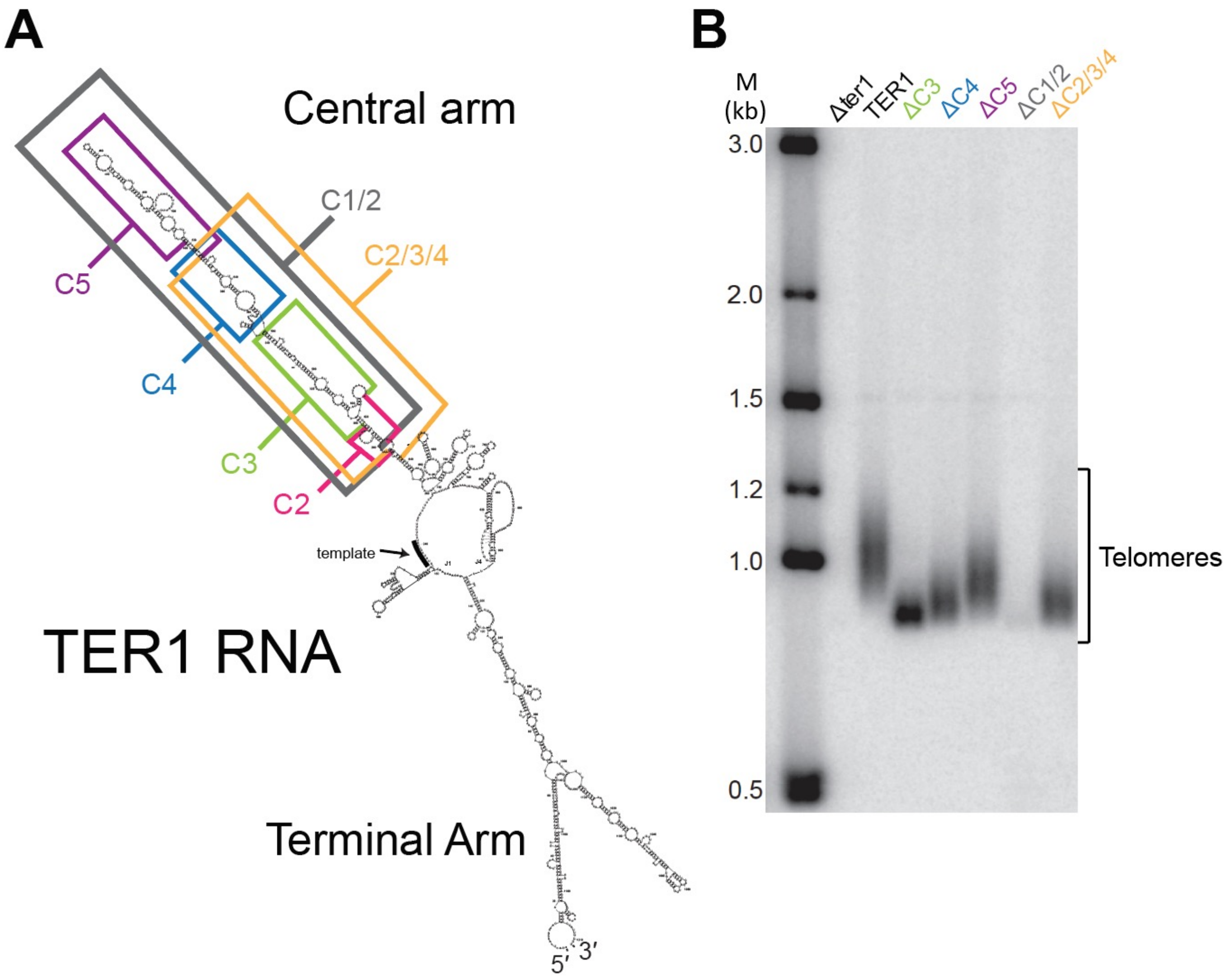


**Supplementary Figure 5.**

**Telomere maintenance in *S. pombe* cells with TER1 Central-Arm truncations.**

**A. Schematic of Central-Arm truncations cloned and tested for complementation *in vivo*.** Boxes indicate the sections of the Central Arm removed and expressed from a plasmid in *S. pombe ter1*Δ cells.

**B. Truncations tested in the Central Arm lead to shorter telomeres.** Telomere Southern blot showing telomeres of the indicated alleles after passaging cells.

### Supp. Fig. 6

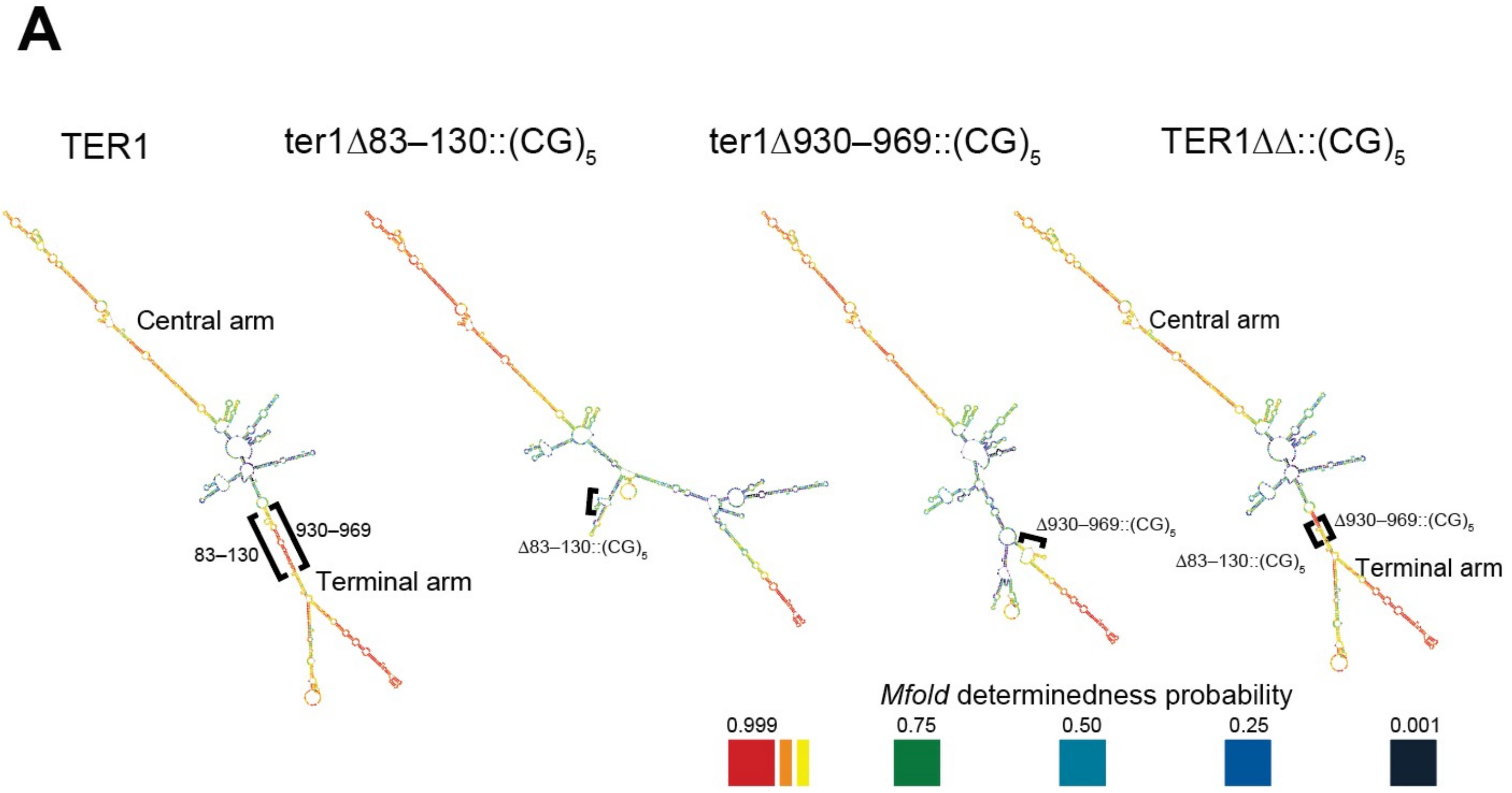


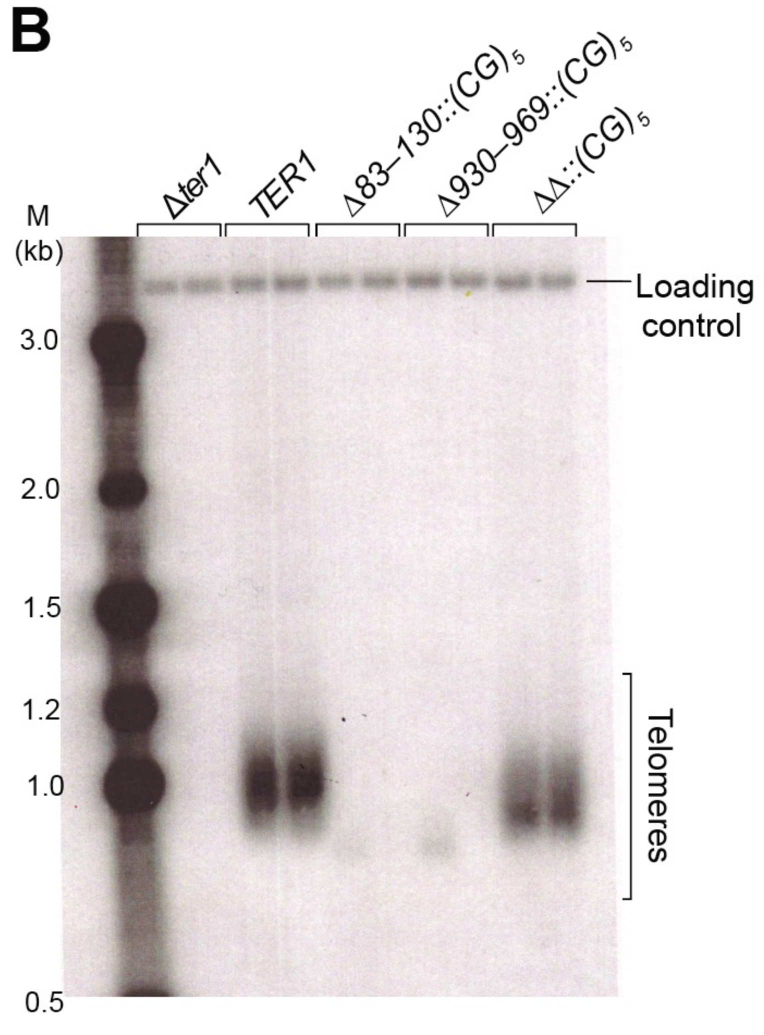


**Supplementary Figure 6**.

**Experimental evidence for the Terminal Arm: compensatory base-substitution mutants on each side of the predicted arm allow function *in vivo* when combined.**

**A.** The nucleotides substituted with $(CG)_5$ on each side of the Terminal Arm are shown on the secondary structures. Since phylogenetic information (covarying base pairs in evolution) are not appropriate to use for modeling of *mutant* TER1 RNAs, bioinformatic lowest-free-energy modeling was employed to compare predicted folding of mutant RNAs compared to wild type. Thus, the models shown in **A** were determined using solely *Mfold*. (Note: this software does not predict pseudoknots.) The rainbow color spectrum in the folds shown is from the output of the *p-num* (or *SIR_GRAPH*) feature of *Mfold* (Zuker and Jacobson, *RNA* 1998). The red end of the spectrum has energetically higher-confidence, or “determinedness,” of the folding prediction based on *Mfold*’s lowest-free-energy calculation algorithm; i.e., the high-determinedness folded conformations also appear in slightly less-negative-$\Delta G$ calculations. Note that the overall folding of TER1 is predicted to be largely restored in the ΔΔ mutant compared to the single Δ alleles.

**B.** Telomeres are lost in single mutants, whereas they are restored in the double-mutant allele to near wild-type length, strongly suggesting these regions are paired in the wild-type TER1 telomerase RNA.

Supp. Fig. 7

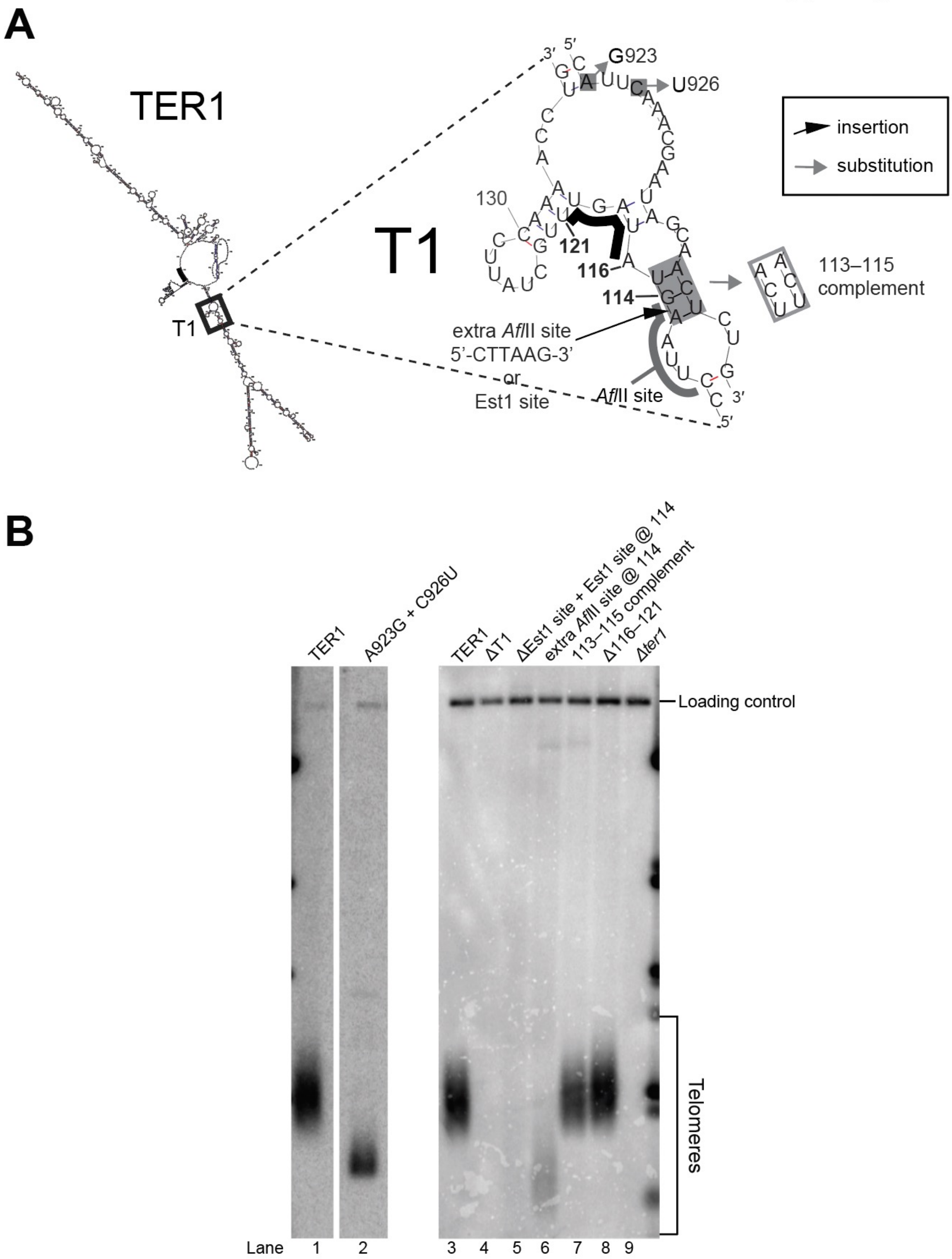


**Supplementary Figure 7**

**Mutational analysis of the essential T1 region of TER1.**

**A. Schematic of the mutants generated in the T1 region.** Mutants designed and tested *in vivo* include deletion (ΔT1), truncations (Δ116–121), nucleotide substitutions (A923G+C926U, insertions (extra *Afl*II site@114), and a predicted base-pair substitution (113–115 complement).

**B. Telomere Southern blot of T1 element-dissecting mutants.**

### Supplementary Table 1

**Base pairs used to constrain *Mfold model based on phylogenetic information*, etc.**

Numbering is based on *S. pombe* TER1 RNA nts.

List of base pairs forced in *Mfold*. *RNAalifold* was used to determine likely-covarying base pairs. These, along with base pairing data from Box *et al.* 2008, were used to constrain the *Mfold* to determine the lowest free-energy structure that also includes all the base pairs listed. Numbering is based on the 1212-nt *S. pombe* TER1.

**Primary sequence order**

| 5' nt | 3'nt |
|---|---|
| 42 | 1165 |
| 51 | 1156 |
| 53 | 1154 |
| 55 | 1151 |
| 61 | 1145 |
| 69 | 997 |
| 73 | 993 |
| 91 | 977 |
| 92 | 976 |
| 117 | 935 |
| 118 | 934 |
| 150 | 235 |
| 153 | 232 |
| 163 | 229 |
| 164 | 228 |
| 165 | 227 |
| 166 | 226 |
| 172 | 191 |
| 208 | 220 |
| 209 | 219 |
| 211 | 217 |
| 260 | 737 |
| 264 | 651 |
| 296 | 623 |
| 327 | 578 |
| 330 | 575 |
| 340 | 561 |
| 341 | 560 |
| 366 | 553 |
| 376 | 544 |
| 377 | 543 |
| 390 | 534 |
| 391 | 533 |
| 402 | 518 |
| 407 | 513 |
| 408 | 512 |
| 415 | 507 |
| 419 | 487 |
| 426 | 480 |
| 427 | 479 |
| 428 | 478 |
| 436 | 469 |
| 438 | 468 |
| 449 | 460 |
| 451 | 458 |
| 656 | 691 |
| 696 | 733 |
| 711 | 718 |
| 745 | 784 |
| 746 | 783 |
| 757 | 769 |
| 834 | 905 |
| 836 | 903 |
| 845 | 895 |
| 1003 | 1135 |
| 1043 | 1093 |
| 1047 | 1089 |
| 1050 | 1084 |
| 1070 | 1075 |

**Rank order**

| 5' nt | 3' nt |
|---|---|
| 118 | 934 |
| 711 | 718 |
| 117 | 935 |
| 208 | 220 |
| 407 | 513 |
| 172 | 191 |
| 408 | 512 |
| 92 | 976 |
| 69 | 997 |
| 426 | 480 |
| 757 | 769 |
| 341 | 560 |
| 73 | 993 |
| 209 | 219 |
| 391 | 533 |
| 211 | 217 |
| 330 | 575 |
| 340 | 561 |
| 377 | 543 |
| 419 | 487 |
| 53 | 1154 |
| 428 | 478 |
| 438 | 468 |
| 1050 | 1084 |
| 746 | 783 |
| 656 | 691 |
| 451 | 458 |
| 1003 | 1135 |
| 376 | 544 |
| 260 | 737 |
| 264 | 651 |
| 1047 | 1089 |
| 427 | 479 |
| 390 | 534 |
| 402 | 518 |
| 327 | 578 |
| 1043 | 1093 |
| 745 | 784 |
| 836 | 903 |
| 845 | 895 |
| 449 | 460 |
| 55 | 1151 |
| 696 | 733 |
| 51 | 1156 |
| 42 | 1165 |
| 415 | 507 |
| 1070 | 1075 |
| 834 | 905 |
| 436 | 469 |
| 366 | 553 |
| 61 | 1145 |
| 91 | 977 |
| 150 | 235 |
| 153 | 232 |
| 296 | 623 |
| TBE bps from Box et al 2008: | |
| 163 | 229 |
| 164 | 228 |
| 165 | 227 |
| 166 | 226 |

#### Supplementary Table 2

| Allele | Plasmid | Nucleotides deleted (truncation alleles) | Total # of nts deleted (truncation alleles) |
|---|---|---|---|
| **WT Ter1** | pDZ995 | | |
| **Ter1 core (528)** | pDZ983 | | |
| **Micro-Ter1 (623)** | pDZ984 | | |
| **ΔC1** | pDZ516 | 298-612 | 315 |
| **ΔC2** | pDZ517 | 276-297, 613-630 | 40 |
| **ΔC1/2** | pDZ321 | 276-630 | 355 |
| **ΔC3** | pDZ318 | 298-343, 558-612 | 101 |
| **ΔC4** | pDZ319 | 345-396, 524-556 | 85 |
| **ΔC5** | pDZ320 | 402-525 | 124 |
| **ΔC2/3/4** | pDZ322 | 277-394, 524-637 | 232 |
| **ΔT1** | pDZ510 | 108-134, 924-945 | 49 |
| **ΔT2** | pDZ515 | 47-64, 1142-1206 | 83 |
| **ΔT3** | pDZ514 | 988-1002, 1127-1141 | 30 |
| **ΔT4** | pDZ513 | 1003-1035, 1096-1126 | 64 |
| **ΔT5** | pDZ512 | 1036-1095 | 60 |
| **ΔT6** | pDZ325 | | |
| **T5@348** | pDZ985 | | |
| **T5@185** | pDZ986 | | |
| **T6@348** | pDZ675 | | |
| **Rev comp T5@348** | pDZ987 | | |
| **WT + T5@348** | pDZ988 | | |
| **WT + T5@185** | pDZ989 | | |
| **83–130::(CG)5** | | 83-130 | 48 |
| **930–969::(CG)5** | | 930-969 | 40 |
| **83–130 + 930–969::(CG)5** | | 83-130, 930-969 | 88 |
| **T1 113–115 complement** | pDZ991 | | |
| **T1 Δ116–121** | pDZ990 | 116-121 | 6 |
| **Est1@109** | pDZ676 | | |
| **T1 point mutants 923 + 926** | pDZ992 | | |
| **T1 w/ extra *Afl* II site** | pDZ993 | | |
| **Vector** | pDZ994 | | |
| | | | Total dispensable nts: 557 |
